# Extended applicability of causal inference to compositional data by reciprocal logarithmic ratio transformation

**DOI:** 10.1101/2021.01.25.428037

**Authors:** Daiki Kumakura, Ryo Yamaguchi, Shinji Nakaoka

## Abstract

Understanding state transition caused by several intrinsic and extrinsic factors such as environmental change and increased stress of human activities has been a significant focus in ecological studies. Analysis of time-series data is indispensable to identify a causal relationship between the possible factors and community change. Among several time-series analysis, a nonlinear time series analysis method called empirical dynamic modeling (EDM) has been recently applied to infer causality of community change that arose from intraspecies interactions. EDM allows model-free analysis to estimate the degree of action strength of intraspecific and interspecific interactions at the population level. Convergent cross-mapping (CCM) is an empirical dynamic analysis method used to suggest the existence of causality by reconstructing the state space of a dynamical system from a time series of observations without assuming any explicit mathematical equations. Although CCM allows for inferring directional interpretation of causal relationships from multivariate time series data, one of the major challenges is its non-applicability to compositional data, a common representation of next generation sequencing data such as microbiome. This study aimed to explore a practical approach applied explicitly to compositional data analysis. More specifically, we propose a heuristic but useful transformation that enables CCM to be applied to compositional data. The proposed transformation has demonstrated its applicability to compositional data equivalent to the conventional CCM to untransformed data. Application of the proposed transformation to sequence-based microbial community profiling data provides reasonable implication to the possible causal relationship during state transition.

**Author summary:** Several types of ecological studies such as trophic changes in lakes, marine plankton communities, forest ecosystems, terrestrial ecosystems, and interactions between plants and soils have employed time-series analyses for identifying the factors that might cause state changes. With the rise of next-generation sequencing, our understanding of these ecosystems is expanding to analyze sequence data. A nonlinear time series analysis method, termed empirical dynamic modeling, has been recently applied for analyzing time-series data. Among different empirical dynamic modeling methods, convergent cross-mapping (CCM) is frequently used to infer a causal relationship. Although CCM enables the directional interpretation of causal relationships, it cannot be applied to compositional data analysis. This study proposes a novel type of transformation, Reciprocal Logarithmic Ratio (RLR) transformation, that enables CCM to be applied to compositional data. With RLR-transformation, CCM results for compositional data are comparable to those for absolute data, and it is confirmed that the transformation is applicable to sequence data as well. The RLR-transformation is expected to provide a better understanding of ecological interactions by estimating causal relationships in compositional data.

## Introduction

Various types of ecological studies, such as trophic changes in lakes [1], marine plankton communities [2], forest ecosystems [3], terrestrial ecosystems [4], and interactions between plants and soils [5], have performed time-series analyses, with particular focus on the analysis of identifying state transition using time series data. These studies have mainly focused on determining the factors that may cause state changes. In such studies, network analysis has been extensively used to improve the understanding of interactions among organisms or between organisms and their environments. For example, some studies have focused on the distribution of barnacles in reef intertidal zones [6], predator-prey relationships between fish and plankton [7], the interaction between trophic and non-trophic interactions affecting biological communities [8], and the network structure between plants and bacteria [9]. On the other hand, existing network analysis methods are less applied to infer causal relationships because of the following reasons. First, undirected graphs, as a typical representation of interaction among elements restrict in network analysis, identify the cause and result of the relationship among elements. Second, most interactions are reconstructed from co-occurrence observations, essentially unidirectional without considering any intervention. Third, when analyzing real-time data of complex ecosystems, determining the main inducers of ecological changes and the direction of the interactions between factors is difficult without using temporal network analysis methods [10].

Recently, a nonlinear time series analysis method called empirical dynamic modeling (EDM) has been applied for analyzing time series data [11]. EDM has several characteristic features: It determines the complexity (dimensionality) of a system, distinguishing nonlinear dynamical systems from linear stochastic systems, and quantifying nonlinearity (i.e., state dependence), determining causal variables, forecasting, tracking the strength and sign of interaction, and exploring the scenario of external perturbation [12]. EDM also allows model-free analysis of community changes to estimate the degree of action strength of intraspecific and interspecific interactions at the population level. Among different types of EDM methods, convergent cross-mapping (CCM) provides inference for causal relationships by reconstructing the state space of a system from time-series data without assuming any mathematical equations. Specifically, CCM reconstructs an attractor of a dynamical system by applying Takens’ embedding theorem to time series data that are assumed to be generated from a nonlinear dynamical system [13]. The causal relationships (i.e., changing one leads to a change in the other) are analogized between the attractors reconstructed from more than two variables of multivariable time series data [14]. Although CCM enables directional interpretation of causal relationships between series, users face several difficulties. For example, its inapplicability to compositional data analysis is a major limitation.

Microbial community profiling based on sequence data uses compositional data. Here we explain why careful consideration is required to infer directionality between a couple of compositional time-series datasets. An intuitive explanation is as follows. This may be because *x*_1_ and *x*_2_ were *x*_1_ and *A* − *x*_1_(*A*: *constant*) relationships, called the constant sum constraint, so in the case of the relationship *x*_1_ → *x*_2_, the constant sum constraint causes the information in *x*_2_ to contain *x*_1_, even though the information in *x*_2_ would not normally be contained in the information in *x*_1_. As a result, opposite directionality is generated by compositions when it should be unidirectional; hence, there is a possibility of false detection. In this study, we empirically show that CCM is still applied to compositional data is if constant sum constraint is appropriately released.

In this study, we focus on transformation and scaling, which give rise to a new opportunity to extend the applicability of the CCM method to compositional data. More specifically, we propose a new transformation and scaling method, that we call Reciprocal Logarithmic Ratio (RLR) transformation. The organization of the present paper is as follows. In the next section, as a validation, we show that the CCM with RLR-transformation for compositional data gives comparable results to those of the conventional CCM for absolute data. Furthermore, we demonstrate that the CCM with RLR transformation can be applied to sequence data that are commonly compositional. Based on the validity of our proposed approach, we apply RLR transformation to a microbiome dataset and obtain biological implications for causal relationship of microbial species interactions.

## Results

In the case of community structure analysis using 16S rRNA genes, the data output by conventional next-generation sequencers sample only a part of the community, and therefore cannot deal with the absolute amount of species number in the community. In addition, primers are used during sequencing to amplify nucleic acids to enable the detection of DNA fragments, so when comparing the obtained samples, the relative amounts should be compared. Hence, sequence data output should be often analyzed as compositional data, lending to the possibility of detecting false causal relationships and interactions by CCM as intuitively explained above. Considering the possibility of false detection, the conventional CCM can only be applied to count data perse, limiting its application scope. There exists a reasonable approach to uncover the limitation. For example, sequence data can be weighted by the number of copies of DNA. More specifically, the use of digital PCR allows for absolute quantification [15]. However, this method is in general costed to implement in every sequencing, and it is still difficult to analyze data that are not subjected to this quantification method. Therefore, since the conventional CCM cannot be applied to compositional data, including sequence data, the current scope of CCM application is restricted to estimate causal relationships only for absolute count data.

In this study, CCM was applied to data that was subject to various transformations. The centered log-ratio (clr) transformation introduced by Aitchison [16] is widely used in various fields [17–25]. The Reciprocal Logarithmic Ratio (RLR) transformation is a novel transformation proposed in this study. Assume that the compositional data consist of two variables, denoted by *X* and *Y*. This transformation transforms and scales *X* and *Y* data (*X* + *Y* = 1) into 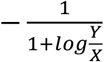 and 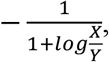, respectively. For these transformed variables, results of CCM were compared based on Cross Map Skill (CMS) and *ω*. CMS is the correlation coefficient between the predicted value and measured value of the other, while *ω* is the difference of CMS. The RLR transformation is tested on variable types of datasets: reference synthetic data and reference real datasets are used for validation, while a sequence dataset is used to demonstrate the references of the RLR transform to infer a causal relationship. A model system of nonlinear dynamics was employed as a source of generating reference synthetic data (see Methods S1 for details). We also examine the effects of changing parameters on reference synthetic data to examine to validate the RLR with reference real datasets; we used the datasets introduced in the CCM proposal paper [14]. Finally, to apply to an insensitivity (robustness) of obtained results a real dataset of sequence dataset, we used the microbial community data [26].

### Evaluation for the applicability of CCM on the reference synthetic data generated from causality enable model under various transformation

To test whether CCM can be applied to compositional data as well as absolute RLR-transformed absolute data, causal relationships were estimated by CCM on reference synthetic data generated from *x*_1_ → *x*_2_ model (causality enable model) under several transformations. Note that a positive value for *ω* implies a possible directional causality from *x*_1_ and *x*_2_. It was found that CCM can be used for compositional data if we apply RLR-transformation. Fig 1 shows the changes of *ω* with respect to the data points. Labels include “Absolute,” “Log Absolute,” and “Reciprocal Logarithmic Ratio (RLR) Transform.” “Absolute” and “Log Absolute” show a decrease of *ω* as the time point increased. “Composition,” “Log Composition,” and “clr Transform” had *ω* values of zero throughout. Fig 1 (above right-hand corner) shows the average of *ω* values for each data point. Positive *ω* values are found for “Absolute,” “Log Absolute,” and “RLR-Transform.” In contrast, “Composition” and “clr-Transform” values are negative, while “Log Composition” value is slightly positive. For each dataset, the population number changes is shown in Fig S1. “Composition,” “Log Composition,” and “clr-Transform,” and the sum of *x*_1_ and *x*_2_, respectively, are always constant at a given point in time, suggesting the existence of a constant sum constraint. By contrast, no constant sum constraint seems to be found for “Absolute,” “Log Absolute,” and “RLR-Transform.” In conclusion, RLR-transformation can successfully estimate the causal relationship for compositional data as well as for absolute data.

**Fig 1.**
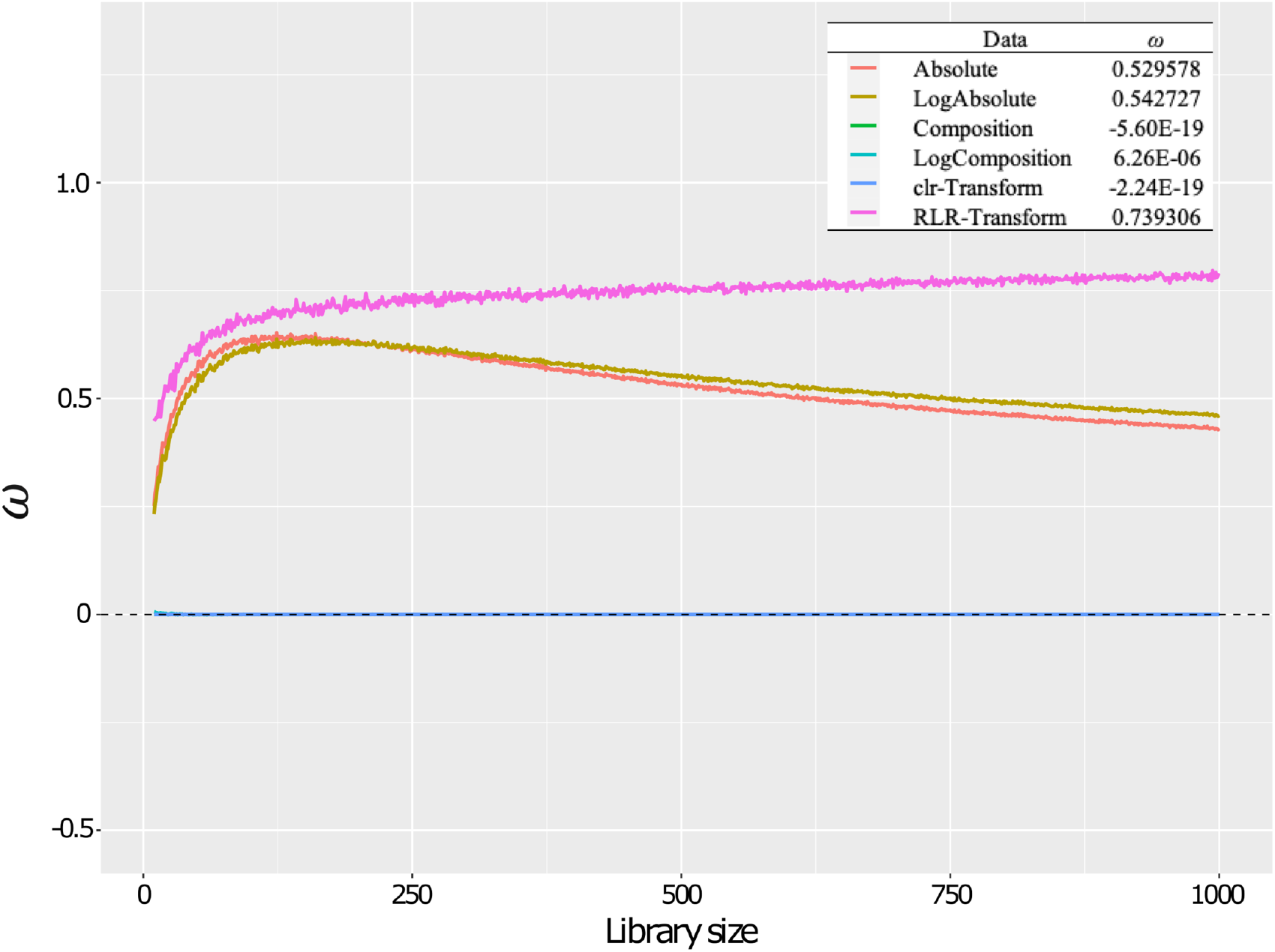
RLR-transformation is an equivalent alternative to the conventional CCM for compositional data. Five transformations were performed using “Absolute” as a reference standard, followed by CCM to each transformed dataset. In addition, the average of the correlation coefficients calculated for each transformation is shown in the upper right-hand corner. Indicators of causal relationship shown that “Absolute,” “Log Absolute,” and “RLR” are positive, while the others are almost zero. The “RLR” exhibits at the same level of “Absolute,” even though the data were converted from “Composition.”

### Dependence of CCM results on parameter

To further investigate whether the RLR transformation can cancel out the inference of constant sum constraint. Parameters were varied to check whether causal relationship can be detected for a wider range of parameter varied, *x*_1_ → *x*_2_ model was used to compare absolute and compositional data with correlation coefficient 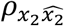. The correlation coefficients between the predicted and measured values of *x*_2_ from *x*_1_ are plotted for each parameter in Fig 2. *x*_2_ is independent of *x*_1_, no correlation should be expected between the predicted and measured values of *x*_2_. However, “Absolute” had positive values for 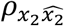, especially for r_2_ ≥ 3.7 & 0 ≤ *β*_21_ ≤ 0.4 & 3.55 ≤ r_2_ ≤ 3.57 where 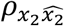 was close to 1. For “Composition,” 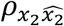 was positive overall. For “RLR-Transform,” 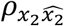 was lower than “Absolute.” 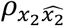 was close to 1 at r_2_ ≥ 3.8 & 0.1 ≤ *β*_21_ ≤ 0.4 & 3.55 ≤ r_2_ ≤ 3.57. The RLR-transformed data showed the same parameter dependence as the absolute data. Although both Absolute and RLR-transformed data are limitations for parameter dependency [27], RLR-transformed data can be applicable to compositional data constant sum constraint of compositional data.

**Fig 2.**
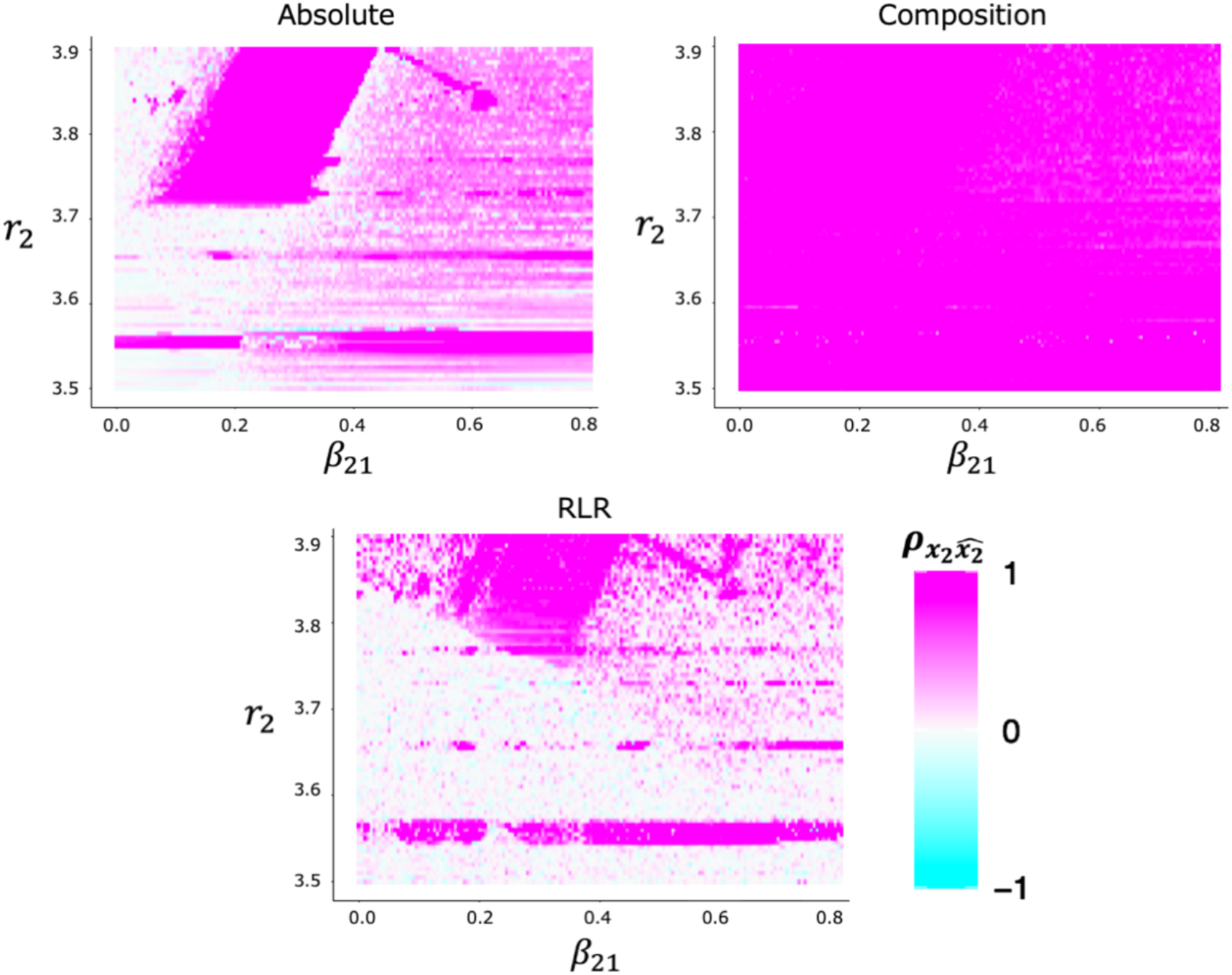
Parameter dependence of each transformation. Each panel represents the correlation coefficients between the observed and predicted values of *x*_2_ from *x*_1_ with respect to time points. The results of “Absolute” show a high sensitivity as the CCM results depend on the parameter choice, as explained by Mønster et al. [27]. On the other hand, “Composition” was unchanged (less sensitive) in almost all cases.

### Testing of real data

Since the effectiveness of the RLR-transformation was confirmed on simulation data, in order to validate the applicability of RLR-transformation on reference real data, we used well known real data as another type of reference for the validation of RLR-transformation. The reference real data have been analyzed by CCM and whose causality is well verified. In the analysis, correlation coefficients (correlation coefficients to CMS) and root mean squared error (RMSE) were calculated as the indicator of accuracy between the predicted value and observed value. The validation results show that the RLR-transformed data give similar results to the absolute data. The results of CMS and RMSE comparisons for the three different datasets are shown in Fig 3. For the experimental results of the predator-prey system, the results of “Absolute” show that the directionality from *Didinium* to *Paramecium* is stronger than that from *Paramecium* to *Didinium* (Fig 3A). For RMSE, the value of the *Didinium* to *Paramecium* direction is lower than that of the opposite direction. For “Composition,” CMS and RMSE shown the same values in both directions and a constant sum constraint is obtained (Fig 3B). For “RLR Transform,” no saturation is observed similar to that observed in “Absolute”, but the direction from *Didinium* to *Paramecium* is stronger than the opposite direction (Fig 3C). Moreover, the *Didinium* to *Paramecium* direction is lower than the opposite direction for RMSE similar to “Absolute.” As another example, the relationship between northern anchovy (*Engraulis mordax*) landings and sea surface temperature (SST) measured at Scripps Pier and Newport Pier in California was also compared in the same way with three datasets (Fig S2). Under the RLR-transformation, similar results to the original data are obtained with CMS and RMSE. In summary, it was confirmed that the RLR-transformation yields the same results as in absolute quantity data, even for real data after compositional transformation.

**Fig 3.**
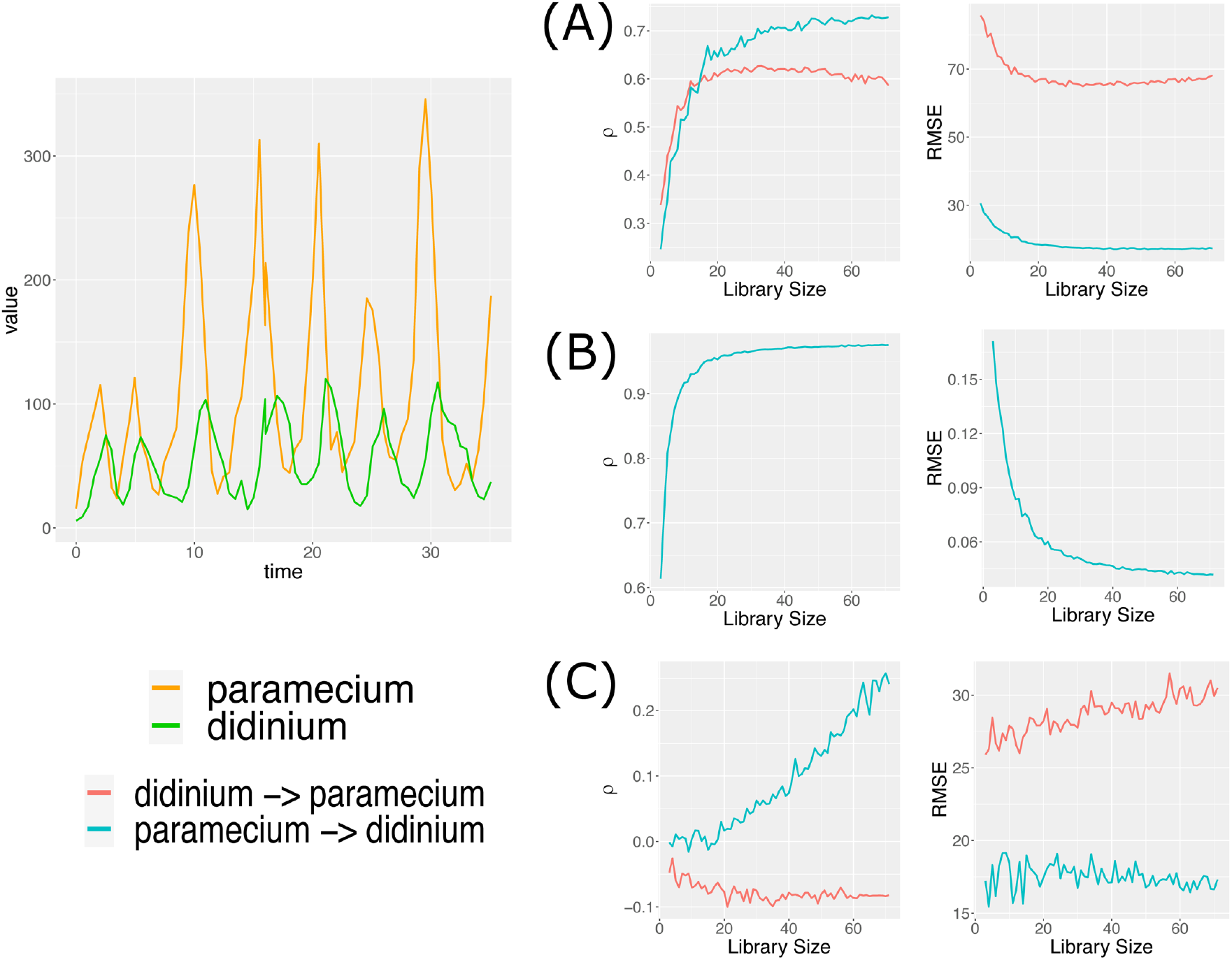
Validation of RLR-transformation on real data. Using “Absolute” as a reference standard, “Composition” shows the same value, and hence, the changes are overlapping. “RLR-Transform” was transformed from the data in “Composition” and showed the same behavior to the results exhibited by “Absolute.”

### Application to real compositional data

Confirmed the effectiveness of RLR-transformation for real data, we then compared CMS and RMSE between operational taxonomic unit (OTU) raw data represented as compositional, and RLR-transformed data in order to validate the applicability to sequence data. The results shown in figure 4 imply that the RLR-transformed data are consistent with the microbe relationships reported in previous studies, while non-transformed data conflict with previous studies. Of note, the relationship between *Clostridium* spp. and *Ruminococcus gnavus* differed before and after the conversion. For *Clostridium* and *Bacteroides* spp., the correlation coefficients are similar, but RMSE shows the opposite changes. This implies that there exists a difference in the CCM results before and after the RLR-transformation. The validation of the RLR-transform is discussed in “Summary and perspectives” in detail.

**Fig 4.**
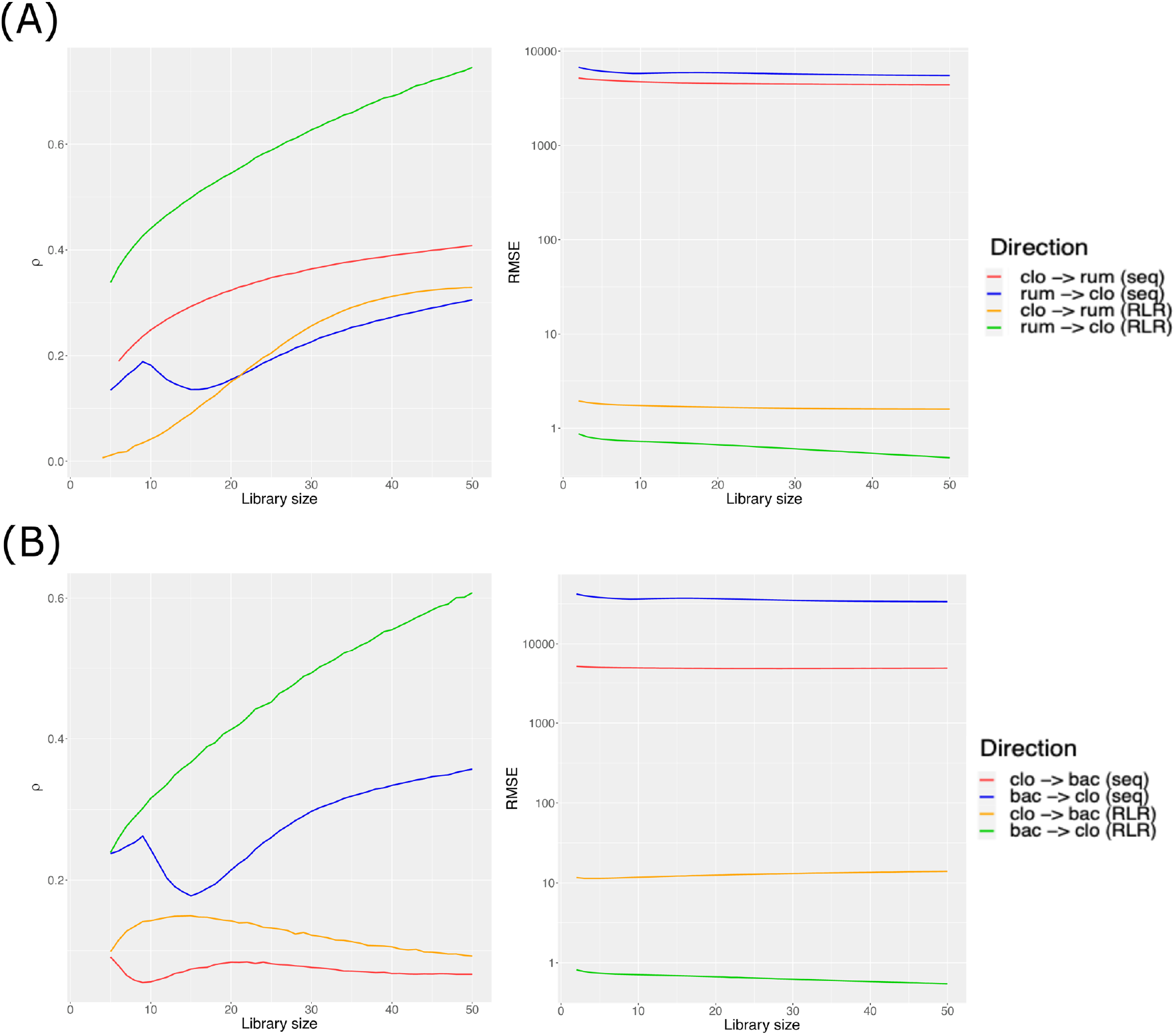
Application of RLR-transformation to sequence data. Time series datasets of 16S rRNA gene sequencing obtained from fecal samples of an infant are compared with the inferred causal relationship by CCM on the OTU count and after RLR-transformed data [26]. (A) shows the relationship between *Clostridium* spp. and *Ruminococcus gnavus*, and (B) shows the relationship between *Clostridium* and *Bacteroides* spp. The red and blue lines represent the OTU count data, and the orange and green lines represent the RLR-transformed data used to calculate the CCM. All known relationships are in good agreement with the inferred results by CCM for the RLR-transformed data.

## Discussion

In this study, we examined transformation and scaling can extend the applicability of CCM for compositional data. The proposed method, RLR-transformation has demonstrated its applicability to broader the score of the CCM. In fact, using *x*_1_ → *x*_2_ model, the effectiveness of the RLR-transformation was verified by comparing it with various transformations, and by comparing the dependence on parameters, initial values and variables, and by analyzing the real data. With the RLR-transformation, CCM infers an appropriate causal relationship for compositional data similar to those for the raw data without any transformation. In the following subsections, we discuss the inferred causality of CCM for various data and elements as well as the newly proposed RLR-transformation.

### Different conversions versus detectability

For “Composition” and “clr-Transform,” the sum of *x*_1_ and *x*_2_ at a given point was always the same, indicating the existence of a constant sum constraint (Fig S1). The constant sum constraint refers to the fact that if the percentage of the D-1 variable is determined when the number of variables in the compositional data is set to D, the percentage of the remaining variables is uniformly determined by subtracting them from the sum of the variables. Compositional data have only one less actual degree of freedom or dimension for the number of variables and is arranged in a D-1 dimensional space [16, 28–29]. For “Absolute” and “RLR Transform” there is no constant sum constraint, and hence, no inter-dependent relationship between *x*_1_ and *x*_2_ is not *x*_1_ and *A* − *x*_1_(*A*: *constant*) as seen in the result (Fig 1). Therefore, it is suggested that the RLR-transformation released the compositional data from the constant sum constraint and yielded the same CCM results as the absolute data.

### Parameters dependence

Since the reference synthetic model does not include *x*_2_’s information in *x*_1_. Therefore, the prediction index of the reverse causality *x*_2_ → *x*_1_ should be 0 in ideal, and the correlation coefficient between the predicted value of *x*_2_ implicated from the data for *x*_1_ and the observed value must be close to zero. However, depending on the parameters, 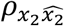 as a predictive index changed (Fig 2). In particular, the constant sum constraint inherent to “Composition” resulted in a high correlation even for reverse causality. In fact, Mønster and Sugihara et al. [14, 27] have already reported that CCM does not correctly predict the direction of causality when the coupling is so strong that it results in the synchronization of variables. For the DE-transformed data, it was confirmed that the prediction accuracy was improved to the level of absolute data. This may be due to the fact that the RLR-transformation unlocks the transformed data from the constant sum constraint. In summary, CCM with RLR-transformation is capable of detecting directional causality for compositional data.

### Verification of the effectiveness of “RLR-Transformation” on reference real data

In the real data analysis, the correlation coefficient and RMSE are used as two predictive indices of CCM to test the effectiveness of the RLR-transformation. Fig 3 shows that the directionality from *Didinium* to *Paramecium* is stronger than the opposite direction in “Absolute,” which is similar to the result reported by Sugihara [14]. For “Composition,” it is probable that the relationship between *Paramecium* and *Didinium* is interdependent because of the constant sum constraint. However, the results for “RLR-Transform” are similar to those of “Absolute,” although there is some variation in each library size, and the graphs for RMSE are similar to those of “Absolute.” Together with, the results of Fig 3 and Fig S2 suggest that the RLR-transformation can be used to apply CCM to compositional data.

### Application of “RLR-Transform” to real data

We apply RLR-transformation to microbiome data to infer possible causal relationships among bacterial species in [26]. Fecal samples, collected weekly between 5.5 and 17 months old of a female infant who was an asymptomatic carrier of *Clostridium* spp., were analyzed by 16S rRNA gene sequencing. From the results of Fig 4, after the RLR-transformation of *Clostridium* spp. and *Ruminococcus gnavus*, it is speculated that *Ruminococcus gnavus* is the cause and *Clostridium* spp. is the result of this relationship. In fact, *Ruminococcus gnavus* is known to convert lithocholate to ursodeoxycholate and plays a major role in the formation of ursodeoxycholate in the colon [30]. Ursodeoxycholate is known to inhibit spore formation [31]. An increase in *Ruminococcus gnavus* is known to contribute to the consequent loss of *Clostridium* spp., which is consistent with the present results [26]. For *Clostridium* and *Bacteroides* spp., *Bacteroides* spp. was the cause, and *Clostridium* spp. was the result. The gut environment is known to form a barrier to *Clostridium* spp. colony formation by competing for metabolic substrates and producing inhibitors of *Clostridium* spp. [32]. For example, the relative abundances of *Clostridium* and *Bacteroides* spp. are inversely correlated in the human gut [33]. *Bacteroides* spp. have been shown to form a resistance barrier to *Clostridium* spp. by competing for monosaccharides such as glucose, N-acetylglucosamine, and succinate and sialic acids [34–38]. Previous studies have shown that an increase in *Bacteroides* spp. contributes to a decrease in *Clostridium* spp., which is consistent with the results of the present study.

### Summary and perspectives

We demonstrated that an appropriate transform recovers the correct inference of causality even for compositional data. The effectiveness of the RLR-transformation was confirmed through the validation on synthetic and real data. The analysis of sequence data showed that the RLR-transformation can be used to estimate the interactions and causal relationships in a microbial community profiling which is often represented as compositional data. RLR-transformation used in this study is expected to produce results similar to those of absolute count data without causing false positives. The approach presented in this paper can open an opportunity for CCM and other interaction estimation analysis to be widely available for sequence data using NGS.

## Materials and Methods

### Model system

The logistic map has long been employed as a standard model system of nonlinear dynamics, exhibiting regular periodic behavior as well as deterministic chaos [39]. In particular, generalization of the logistic map allows representing population dynamics of multiple species.

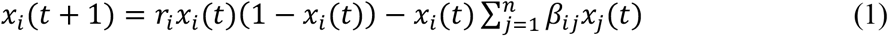

Note that interspecific competition is represented by *β*_*ij*_*x*_*i*_(*t*)*x*_*j*_(*t*), where *β*_*ij*_ is the strength of competition (*i*, *j* = 1,2, …, *N*).

### RLR transformation

Reciprocal Logarithmic Ratio (RLR) transformation is defined as a kind of softmax function. Let the compositional data be *X* and *Y* with absolute data as *x* and *y*, and let us consider the transformation of *X*. Now we assume that both *X* and *Y* have an effect on each other. The logarithmic odds for the likelihood of *X* to *Y* are given by 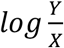. If *X* is the cause and *Y* is the result, the log odds represent the likelihood of the result, and if *X* is affecting *Y*, the odds are high. The log odds are defined as a variant of softmax function, 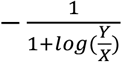, and this transformation is called the Reciprocal Logarithmic Ratio (RLR) transformation. This means that *X* becomes 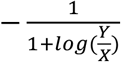 by the RLR transformation.

### Various transformations

In this study, we generated various transformed datasets to apply CCM and compare results. Transformations include “Absolute” (untransformed), “Composition,” “clr-Transform,” and “RLR-Transform.” We also examined logarithmic transformations (to cover more different types of transformation, see Methods S1 for details). For “Composition,” the values at each time point are obtained as a proportion. The “clr-Transform” and “RLR-Transform” are applied to “Composition” (Fig 8). For “clr-Transform,” the centered log-ratio transformation (clr) introduced by Aitchison was used [16]. Clr-transformation maps a composition in the *D*-part Aitchison simplex isometrically to a *D-1* dimensional Euclidean vector. The clr representation of composition *x* = (*x*_1_, …, *x*_D_) is defined as follows:

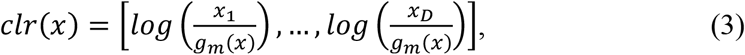

Where 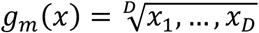.The sum of the component of *clr*(x) is zero [40]. The clr transforms the composition to the Euclidean sample space and hence provides the possibility of using standard unconstrained statistical methods for analyzing compositional data [41]. This conversion method was widely used in geology [17–19]. In recent years, it has been applied in various fields, such as social sciences [20], archaeology[21], agriculture [22], microbiology [23], ecology [24], and evolution [25].

### Convergent cross-mapping

Convergent cross mapping (CCM) is a method for inferring causality in terms of the reconstruction of the state space of a dynamical system from multiple time series data [14]. The basic principle of cross-mapping involves reconstructing system states from two-time series variables and then quantifying the correspondence between them using nearest neighbor forecasting [42]. For two-time series, two attractors are reconstructed by embedding from Takens’ theorem. Then, the effect of one on the other is measured by evaluating the error of the correlation coefficient (Cross Map Skill) between the estimated time series and the actual time series. In this study, we checked the nonlinearity of generated time-series data by S-map which is computed by using R package rEDM, selected the embedding dimension at minimum predictive RMSE, and conducted cross-mapping from the two-time series using the “ccm” function in rEDM. The results were visualized as library size on the horizontal axis and Cross Map Skill (CMS) on the vertical axis. In this study, we introduced a new measure of causality, *ω*, to compare the results of CCM across multiple data sets (Fig 1). Note that *ω* represents the CMS difference between the pair of CCM results for each library size. Using this difference, we compared the changes of CMS by CCM in multiple datasets. More specifically, *ω* is computed as follows:

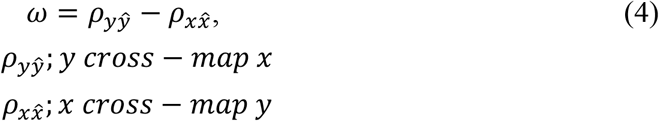

### Validation of CCM using multiple types of data

To validate whether CCM can correctly infer causal relationship, multiple artificial data representing population dynamics of a prey-predator system are generated. First, we considered a two-variable nonlinear deterministic coupled system [14] (see Methods S1 for details). The dataset is transformed “Absolute,” “Composition”, “clr-transform”, and RLR-transform (see S1 for details). CCM was performed for each type of dataset. Then, the directionality was evaluated by *ω*, and compered to examine correct detectability.

### Verification of parameter dependence

We tested whether the results of CCM were affected by parameter values. The conditions for the various parameters and variables were generated according to Mønster et al. 2017 [27] (see Methods S1for details). The data were calculated according to the conditions, with “Absolute,” “Composition,” and “RLR-Transform.” For verification, the correlation coefficient between the predicted and measured values of *x*_2_ was used.

### Validation of the effectiveness of RLR-transformation on reference data

Using the actual data, we validated “RLR Transform” for CCM applicability compared with “Absolute” and “Composition.” For the actual data, we used the two datasets previously employed in [14], and the sequence dataset with fecal microbiota [26]. The first was analyzed using a predator-prey system at the laboratory level, where both the directions of causality exist. This was first studied by Gause in the 1920s and later refined by Veilleux [43], confirming the involvement of *Didinium* (predator) and *Paramecium* (prey) [14]. For these data, CCM was applied, and CMS comparison was performed using the optimal embedding dimension. In addition, to evaluate the accuracy of the predictions, the RMSE values were also compared with each dataset.

### Application to real data

The microbial community profiling data obtained from fecal samples were used to compare differently. Briefly, 16S rRNA gene sequences are obtained from fifty fecal samples, collected weekly between 5.5 and 17 months old of a female infant who was an asymptomatic carrier of *Clostridium* spp. Here, we focused our analysis on the relationship between *Clostridium* spp. and *Ruminococcus gnavus*, *Bifidobacterium bifidum*, and *Bacteroides* spp., which was described in [26]. Microbiome data represented by the operational taxonomy unit (OTU) table were used (see for the detail [44]). Specifically, the time-series data for *Clostridium* spp. and *Ruminococcus gnavus*, *Clostridium* spp., *Bifidobacterium bifidum*, and *Clostridium* spp. and *Bacteroides* spp. were analyzed and compared with the RLR transformed data, and the results were discussed in addition to what has already been discussed in the previous study [26].

## Acknowledgments

This work is supported by JST-Mirai Program Grant Number JPMJMI19B1, the Japan Society for the Promotion of Science (JSPS) Grant-in-Aid (S) 15H0570710 and (B) 18H0266210, and the Ministry of education, culture sports, science and technology-Japan (MEXT) Ambitious Tenure Track program in life science, Hokkaido University.

## Supplementary information

### Methods S1. Detailed methods for each verification point.

#### Comparison of logarithmic transformation with several conversions

In this study, we tested comprehensive transformation with several datasets including compositional data. The data output from the equation is transformed to “Absolute” (no transformation), and “Log (Absolute + 1).” For compositional transformed data “Composition,” we further transformed to obtain “Log (Composition + 1),” centered log-ratio transform “clr-Transform,” and the newly proposed transformation “RLR Transform.” Note that “Log (Value + 1)” and “clr-Transform” are major transformations for converting compositional data.

#### Validation of CCM using multiple types of data

Using the population dynamics model below, we generate 1000 time series datasets of the system with *x*_*i*_(0) = 0.1 as an initial condition. Parameter values are set as follows:

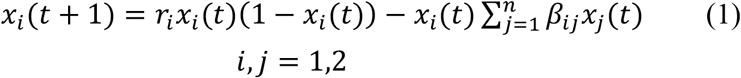

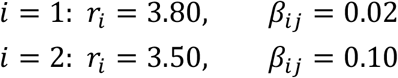

#### Verification of parameter dependence

For system(1), we set a restriction to equations such that *x*_1_ consists of *x*_1_ only, whereas *x*_2_ consists of *x*_1_ and *x*_2_. Synthetic data were generated up to 1400 time points, and 1000 to 1400 time points were used. Parameters and variables are set as follows:

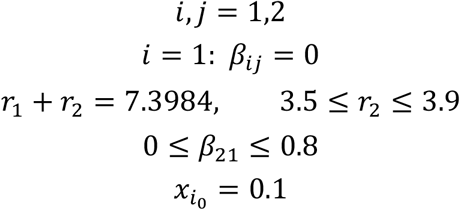

## Appendix S1. Testing of real data on the relationship between anchovy and sea surface temperature.

## Material and Methods

The dataset was analyzed to identify the complex causation in the sardine–anchovy system [14]. The data describe the relationship between northern anchovy (*Engraulis mordax*) landings and sea surface temperature (SST) measured at Scripps Pier and Newport Pier, California. As the data used in this study included negative values, min– max normalization was performed, and the two datasets were normalized. The conversion formula for min–max normalization is given below:

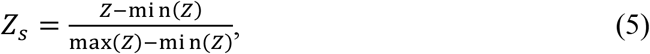

where z is an original value and *Z*_s_ is the normalized value. The raw data were denoted “Raw,” while the compositional data were “Composition,” and the data transformed “Composition” to RLR-transformation were denoted as “RLR-Transform,” respectively. For these data, CCM was applied, and CMS comparison was performed using the optimal embedding dimension. In addition, to evaluate the accuracy of the predictions, RMSE values were also compared with each dataset.

## Result and Discussion

Considering the analysis results of the complex causation in the sardine-anchovy system, the change of CMS for “Raw” are shown in Fig S2 A. This indicates that sea surface temperature affects the landings of anchovies. For “Composition,” CMS and RMSE exhibit the same values in both directions, and a constant sum constraint was obtained (Fig S2 B). For “RLR-Transform,” the changes of CMS and RMSE were similar to that of “Raw,” and the direction from sea surface temperature to anchovy landings had lower values than that in the opposite direction (Fig S2 C). Fig S2 implies that, for “Composition,” the constant sum constraint may be due to the equally strength of 2 variables. For “RLR-Transform,” CMS and RMSE showed the same changes as obtained for “Raw.”

---

**Fig S1.**
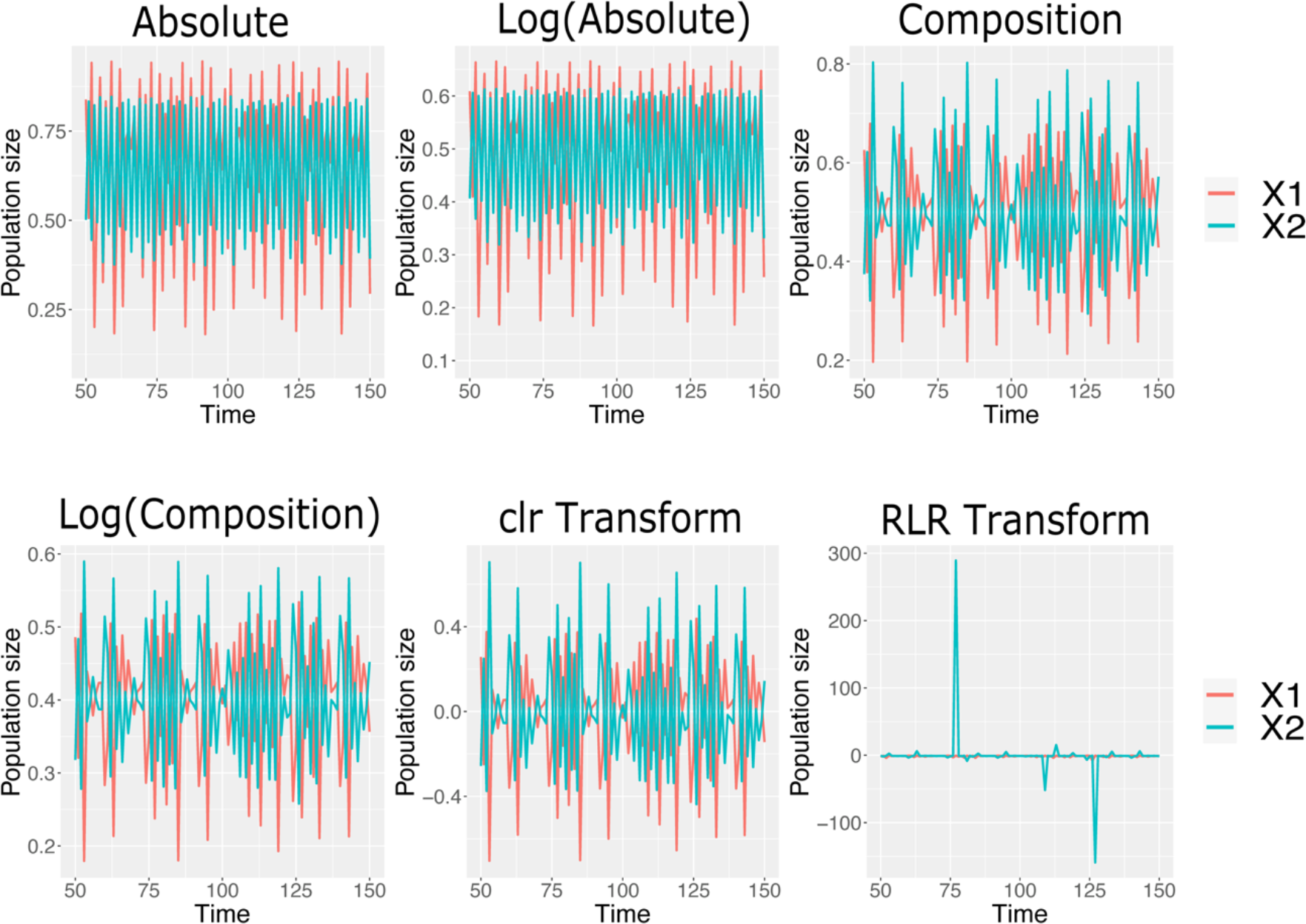
Changes for five transformations with reference to “Absolute.” “Composition,” “Log Composition,” and “clr-Transform” has a constant sum of the two time-series datasets. By contrast, no constant sum is found for “Absolute,” “Log Absolute,” and “RLR-Transform.”

**Fig S2.**
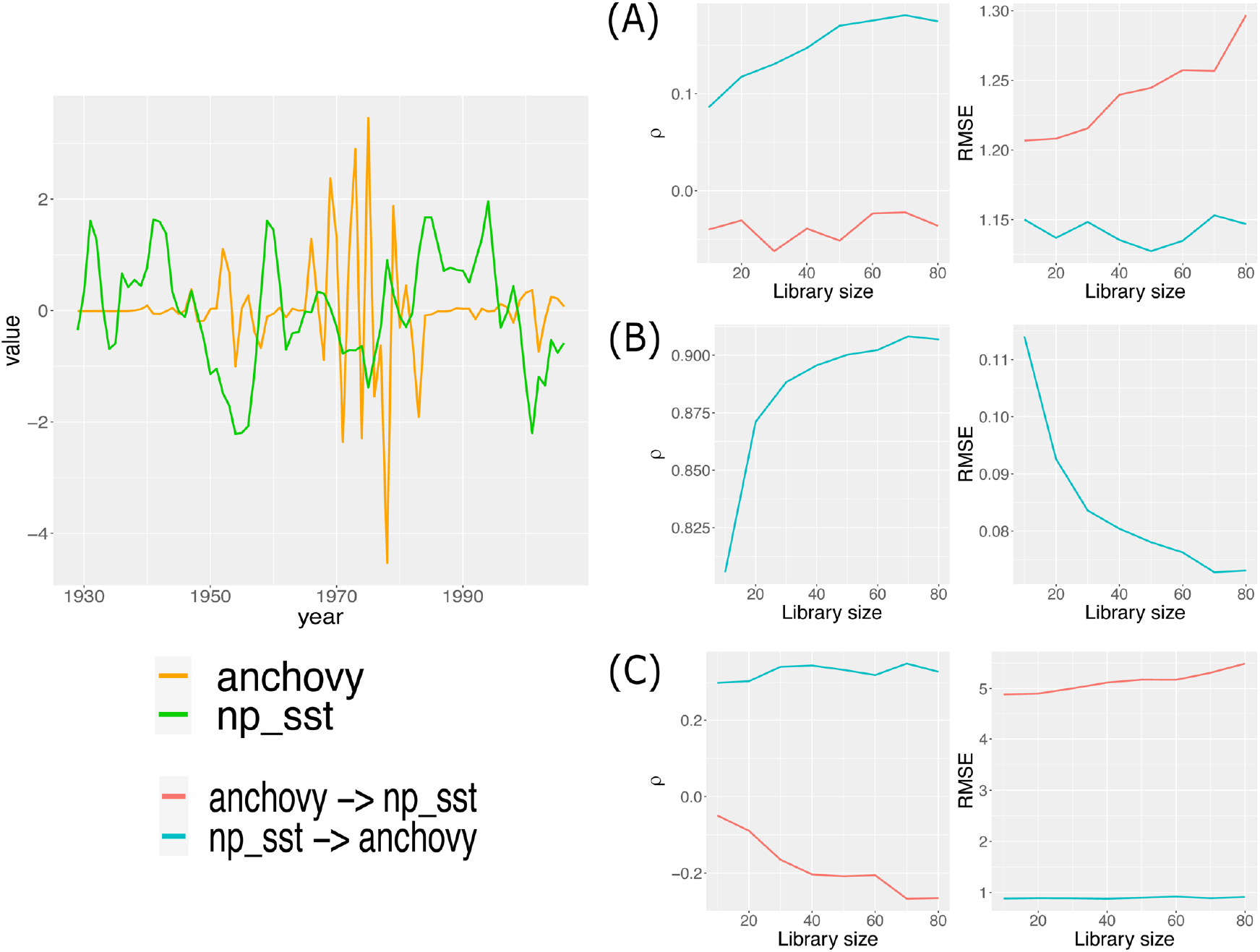
CCM results before and after conversion for determining the relationship between northern anchovy (*Engraulis mordax*) landings and sea surface temperature (SST) measured at Scripps Pier and Newport Pier, California [14]. CCM results for min-max normalized raw data (A), composition (B) and RLR-transform (C). Similar results are obtained for (A) and (C)

**File S1. All the script file used in this study.**

Each R script is used to generate the corresponding figure.

## Notes

https://github.com/DaikiKumakura/RLR_transform

### Competing Interest Statement

The authors have declared no competing interest.

